# Larval zebrafish use olfactory detection of sodium and chloride to avoid salt-water

**DOI:** 10.1101/2020.08.19.258061

**Authors:** Kristian J. Herrera, Thomas Panier, Drago Guggiana-Nilo, Florian Engert

## Abstract

Salinity levels constrain the habitable environment of all aquatic organisms. Zebrafish are freshwater fish that cannot tolerate high salt environments and would, therefore, benefit from neural mechanisms that enable the navigation of salt gradients to avoid high salinity. Yet, zebrafish lack epithelial sodium channels, the primary conduit land animals use to taste sodium. This suggests fish may possess novel, undescribed mechanisms for salt detection. In the present study, we show that zebrafish, indeed, respond to small temporal increases in salt by reorienting more frequently. Further, we use calcium imaging techniques to identify the olfactory system as the primary sense used for salt detection, and we find that a specific subset of olfactory receptor neurons encodes absolute salinity concentrations by detecting monovalent anions and cations. In summary, our study establishes that zebrafish larvae have the ability to navigate, and thus detect salinity gradients, and that this is achieved through previously undescribed sensory mechanisms for salt detection.

## Introduction

All organisms must maintain their internal ionic content within a tight window. However, the mechanisms that animals utilize for osmoregulation are diverse and depend on their environment. Land animals and marine mammals balance the consumption of ion-rich food or liquids with the intake of ion-poor water and excretion of excess ions ^1,2^. On the other hand, fish and amphibians supplement fluid consumption and excretion with the function of a specialized system of epidermal cells, ionocytes, that exchange ions with their surrounding water ^3,4^. Critically, the direction and mechanism of ion exchange depend on the salinity of the animal’s natural environment. Ionocytes of freshwater fish must acquire salts, while those of marine fish excrete them.

Terrestrial animals encounter and balance salt through food intake, leading their gustatory systems to evolve high sensitivity to salt ^5,6,7,8^. They usually use two separate channels for discriminating either appetitive (< 100 mM) or aversive (>150 mM) NaCl concentrations. In vertebrates, these are respectively mediated by epithelial sodium channels ^5^, which specifically allow sodium influx ^9^, and bitter taste receptors ^7^ that are broadly gated by noxious stimulants. By contrast, the primary external salt sensors for fish are unknown at both the tissue and molecular level. However, aquatic animals are directly exposed to changes in environmental salinity, suggesting it might be advantageous for their salt detection to be integrated with a variety of modalities, beyond taste reception, that are suited for environmental navigation, such as olfaction ^10^, somatosensation ^11^, or even the mechanosensory lateral line ^12^. The teleost gustatory system, although clearly implicated in feeding related behaviors, is less likely to play a critical role in salt gradient navigation, since teleosts have lost all homologs of the mammalian epithelial sodium channel family ^13^, and bitter taste receptors do not respond to the relevant salt concentrations ^14^. As such, it is still unclear which molecular mechanisms fish use to detect external NaCl and to appropriately adjust their behavior.

Zebrafish in particular, would benefit from the ability to detect and navigate NaCl gradients since the environments in which they likely evolved are characterized by dramatic changes in local salinity levels: the river basins that surround the Ganges River in India and Bangladesh for example ^15,16^, are characterized by soft, ion-poor water with NaCl concentrations below 1 part per trillion,which can increase locally by orders of magnitude during the dry season ^17^. Importantly, such changes lead to elevated stress and cortisol levels, and are ultimately lethal ^18,19,20^, which makes neural mechanisms for detecting and avoiding salt gradients paramount for survival. Here, we show that zebrafish have evolved behavioral strategies to avoid high-salt environments and that this behavior is mediated by the olfactory system through a subset of olfactory sensory neurons that detect the combined presence of sodium and chloride.

## Results

### Larval zebrafish avoid high salt environments

To test whether larval zebrafish avoid high saline environments, we developed an assay for the detailed observation of larvae swimming in a salt gradient (**1A** and methods). In this assay, four larvae are placed in separate lanes bookended by two agar pads made either from filtered fish water alone (control) or from fish water plus the salt being tested (source). We show that diffusion of sodium chloride (NaCl) from the source generates a persistent and reasonably stable salinity gradient throughout the lanes (**S1A**), and that larvae navigating such a gradient spend significantly more time away from the source than in control conditions where no salt is added (**1B,C**). This aversion emerges alongside the developing salt gradient over the first 15 minutes and is stable afterward (**S1B**). This behavior is robust and demonstrates that larvae as young as 5 days post fertilization (**S1C**) are already equipped with an active strategy to avoid regions of high salinity.

**Figure 1:**
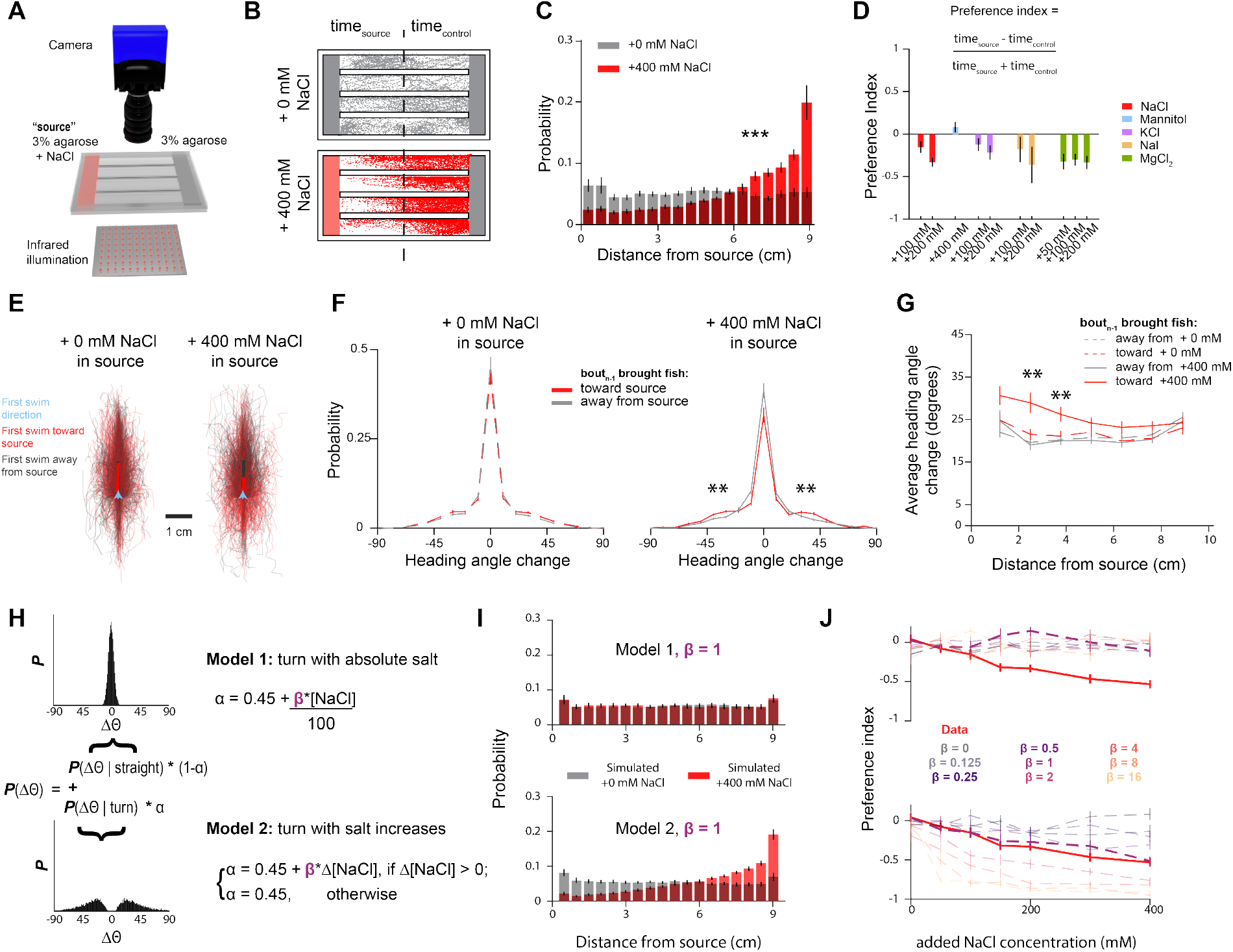
Larval zebrafish avoid high salt environments by responding to increases in salinity. **A.** Schematic of the rig used to perform chemical place preference assays. **B.** Sample experiments when 0 mM (top) or 400 mM (bottom) are added to the source agarose. Individual dots demarcate position of larvae at every 50^th^ frame. **C.** Histogram of positional occupancy by larvae when the source gel contains either 0 mM (n = 30) or 400 mM NaCl (n = 24) added (Mann-Whitney U test p<0.001). **D.** Preference indices toward different cation/anion pairs. **E.** Individual trajectories of 10 bouts that follow a bout that climbs (red) or descends (gray) gradients with 0 (left) and 400 mM (right) NaCl added to the source. Average trajectory indicated by thick lines. **F.** Average turn angle as the larvae swims toward or away from the source as a function of their position in the arena (Bonferroni-corrected t-test, p <0.01). **G.** Difference in average turn angle of bouts that follow an increased salt concentration compared to those that follow a decreased salt concentration for different concentrations of source salt (Bonferroni-corrected t-test, p <0.01). **H.** Description of the two models being simulated: larvae respond to absolute (model 1) or relative (model 2) salt concentrations. **I.** Spatial distribution of larvae in a simulated linear salt gradients when model 1 (top) or model 2 (bottom) are active. **J.** Preference indices toward different concentrations of NaCl that results from simulating fish according to both algorithms for a range of salt sensitivities (*β*).

External sodium chloride fluctuations change several environmental parameters, such as osmolarity, net conductivity, and specific ion identities. Any of these may be detected and used by the larvae to avoid high salt, and in order to dissect the relevance of each parameter, we tested the preference of larvae to a series of compounds that isolate specific factors. For example, a general aversion to increased osmolarity was tested by generating gradients of a sugar alcohol, mannitol, that were equimolar with the previously tested NaCl gradients (**1D**). Under these conditions, the larvae exhibit no place preference. By contrast, zebrafish reliably avoided every ionic solution that we tested, whether it consisted of chloride paired with another monovalent (KCl) or divalent cation (MgCl_2_), or sodium paired with another anion (NaI). These results fail to tell us whether the mechanism underlying NaCl detection is based on a broad conductivity sensor or whether it is specific to different ions, as it is unclear whether or not the mechanisms responsible for avoiding these other ions belong to identical or parallel neural pathways. A detailed resolution of this question necessitates the identification of the relevant sensory neurons, the approach to which is described in later sections.

### Salt avoidance is driven by detecting salt increases

Next, we wished to understand the specific heuristics that larval zebrafish employ to avoid high salt concentrations. We observed that in all conditions, larvae predominantly swim back and forth between the two ends of the arena, while aligned parallel to the longitudinal axis (**S1D,E**). As larvae ascend a salt gradient, however, they reverse directions earlier than during gradient descent (**1E**). This suggests a biased random walk where increasing salt concentrations amplify the larva’s reorientation frequency ^21,22^. To test this hypothesis, we compared the distribution of reorientation angles during bouts that follow a swim toward salt with those that follow a swim away from salt. With no gradient, larvae do not alter their turn statistics between these two cases (**1F**, left). By contrast, during gradient navigation, larvae are significantly more likely to execute a 20-40 degree turn if the previous bout brought them closer to the salt rather than away from it (**1F**, right). Compared to control conditions, turning magnitudes are only affected by the salt concentration as the larva climbs the gradient (**1G**), suggesting salinity increases, rather than absolute concentrations, drive turning.

To test whether a biased random walk is sufficient to explain the larvae’s avoidance behavior, we simulated the ability of virtual larvae utilizing natural swimming and turn statistics to navigate a salt gradient. We compared the performance of these simplified agents in conditions where turning was either upregulated as a function of absolute salinity levels or, alternatively, where turning probability increased after relative increases in salinity (**1H**, see methods). We found that only the latter adequately captured the avoidance behavior of real animals (**1I,J**). A potential concern is that, as formulated, this model predicts that larvae would swim readily toward water with no salt at all, a counterintuitive result given the significantly deleterious effects this would have on the animals (**S1G**). Testing this prediction with deionized water, we found, however, that larvae indeed swim toward the lowest salt concentrations (**S1H**), which suggests that larvae may not seek an optimal external NaCl setpoint, but instead always avoid increasing salinity. A potential explanation for this simplistic, if maladaptive, strategy is that regions of deleteriously low salinity are extremely rare and likely did not impart any selective pressure onto these animals.

We have so far identified that larval zebrafish avoid waters with high salinity, and do so by responding to salt concentration increases. We next wished to identify the sensory modalities that detect salt. However, to identify such sensory regions, we wished to use calcium imaging techniques (*i.e.* light-sheet or two-photon point scanning microscopy) that require us to immobilize the animal’s head. To accommodate this constraint, we designed a stimulation setup (**2A**) to rapidly and reversibly present different chemicals to the larvae’s face (**S2A-C**), while the head and torso are embedded in agarose, and the tail is free to move. We find that larvae in this preparation respond to salt pulses by vigorous tail flicks (**2B**) in a concentration-dependent manner (**2C**). Consistent with the free-swimming behavior, the larvae are most responsive during the onset of a NaCl pulse, corresponding to a recent concentration increase (**2D**). Therefore, we treat this preparation as a reasonable proxy for the more naturalistic condition of a larva swimming freely in a concentration gradient.

**Figure 2:**
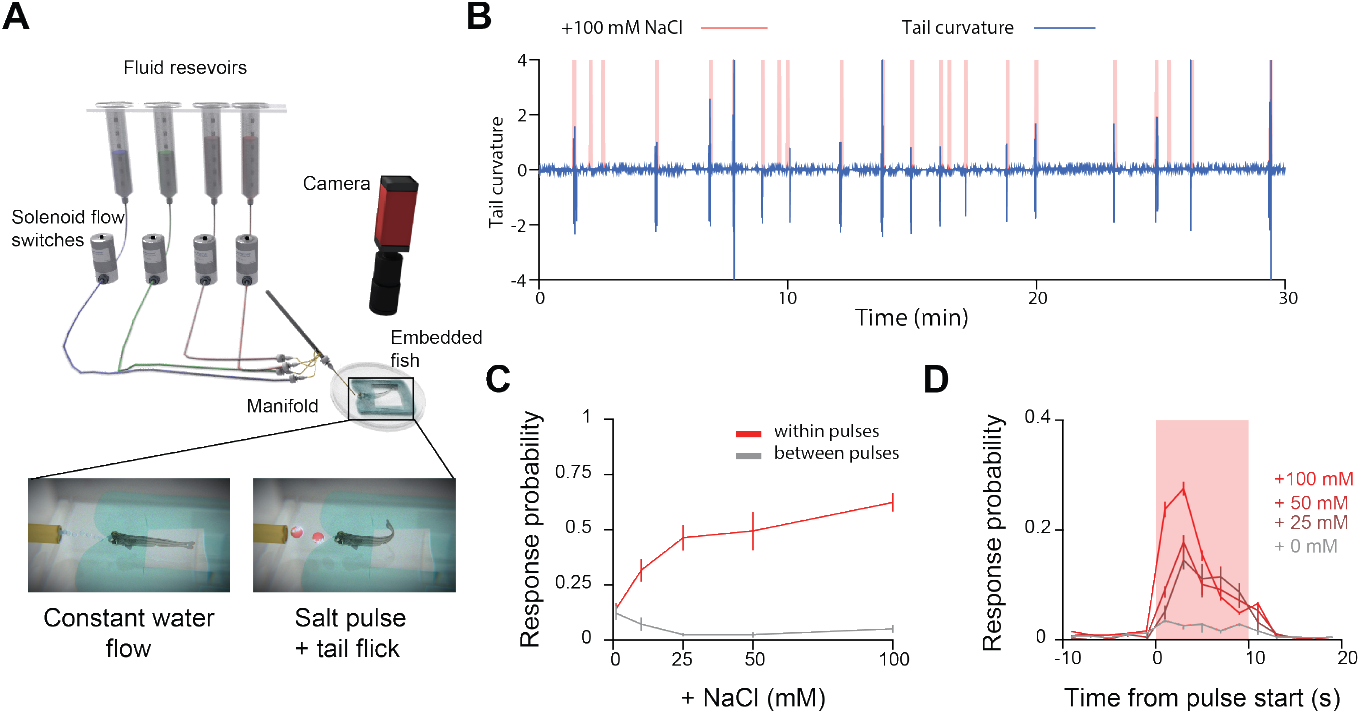
Head-immobilized larvae respond to salt concentration increases. **A.** Schematic of preparation used to stimulate head-embedded larvae. **B.** Sample data from an experiment where a larva is stimulated with 10 second pulses of 100 mM NaCl at random intervals. **C.** Probability that the larva will exhibit a behavioral response to a pulse of a given NaCl concentration (red) and inter-pulse spontaneous behavior rate (gray) to flowing fish water. **D.** Probability of a bout event occurring within two second bins for different concentrations of salt relative to the onset of a pulse.

### Activity in the olfactory and lateral line systems reflects external salinity

To screen for the brain regions most sensitive to such NaCl pulses, we combined our tethered preparation with a custom-built lightsheet microscope (**3A** and **Movie S1**). This allowed us to perform volumetric imaging over most of the larval zebrafish’s brain (**3B**) while delivering pulses of different NaCl concentrations (**3C**) and simultaneously tracking its behavior (Figure 3E). After imaging, we segmented the fluorescence from each plane into activity units (**3D**) using a temporal correlation-based algorithm ^23^. We must note that this algorithm only utilizes correlation across time and incorporates no anatomical features, so these “activity units” may consist of individual cells, neuropil, or combinations of both. To localize segmented units within community standardized anatomical regions, we registered all imaged volumes to the reference coordinate system of the online *Z-brain* atlas ^24^.

**Figure 3:**
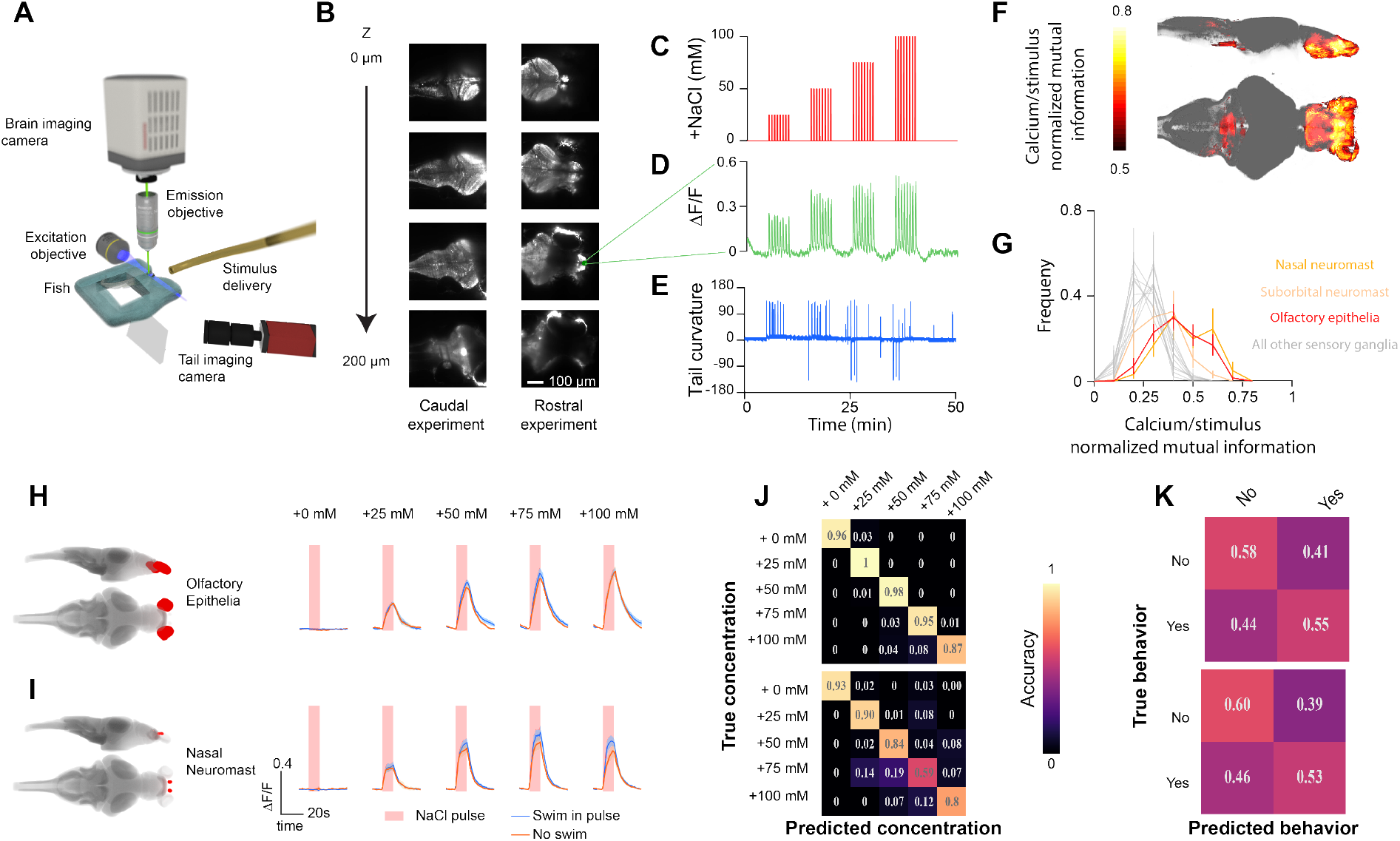
NaCl levels are represented in the olfactory system and and lateral line. **A.** Schematic of the lightsheet microscope. **B.** Sample z-slices taken within a stack. Stacks were collected at 1 Hz. Imaging experiments captured either the rostral 2/3^rds^ or caudal 2/3^rds^ of the fish. **C.** Arrangement of salt pulses during each experiment. Zebrafish experienced escalating concentrations of NaCl pulses in five minute blocks. **D.** Example calcium signal from an activity unit in the olfactory bulb. **E.** Example tail curvature trace during imaging experiment. **F.** Average stimulus correlation at each voxel of the *Z-brain* across 15 fish. **G.** Histogram of normalized mutual information in units from each of the sensory ganglia in the *Z-brain*. **H.** Stimulus triggered responses of top NaCl encoding units from the olfactory epithelia, averaged across fish, and their locations within the *Z-brain*. Responses are separated into trials where the fish swam (blue) or did not swim (purple). Mean SEM across fish. **I.** Stimulus triggered responses of top NaCl encoding units in the nasal neuromast averaged across fish. **J.** Confusion matrices depicting *stimulus* classification accuracy of support vector machines trained from activity in the olfactory epithelia (top) and nasal neuromast (bottom). **K.** Confusion matrices depicting *behavioral response* classification accuracy of support vector machines trained from activity in the olfactory epithelia (top) and nasal neuromast (bottom).

In order to identify any regions that carry information about salinity, we calculated the mutual information between each unit and the delivered salt concentration. Averaging this value across fish for each voxel of the *Z-brain* atlas reveals stereotypic strong NaCl representation within the fish’s olfactory system, which includes the epithelia, bulb, and posterior telencephalon (**3F**) ^25^. Of all sensory ganglia, only the rostral-most neuromasts and olfactory epithelia contain units whose activity reflects NaCl concentration (**3G**). In both of these modalities, we observe concentration-dependent activity that is independent of the animal’s behavioral response (**3H,I**). Furthermore, unlike the behavior, which quickly subsides a few seconds into a pulse, these regions sustained or even ramped their calcium levels throughout the stimulus period. This suggests that the olfactory epithelia and neuromasts of the lateral line contain sensory representations of external salt concentration, and do not, alone, reflect when an animal will respond. To verify this, we trained support vector machines to classify either the stimulus or behavioral response from the activity of units within these regions. As predicted, these classifiers were independently able to predict the stimulus, but failed to predict whether the animal produced a behavioral response (**3J,K**). Thus, the neural substrate for the computations that extracts changes in salt concentration and generates behavioral responses likely occurs somewhere downstream of these primary sensory regions.

Outside of the olfactory system and neuromasts, most regions of the brain share low mutual information with the sensory stimulus, including brainstem regions previously reported to show highly stereotyped responses to tastants ^26^. One exception is a midbrain cluster of units near the dorsal raphe and interpeduncular nucleus (**3F**), whose activity patterns share high mutual information with the animal’s behavior, as well as the combination of stimulus and behavior (**S3A,B**). We therefore propose that these areas represent a substrate for the sensorimotor transformation of salt elevations into action, as supported by the fact that both stimulus and action can be decoded from these regions (**S3E,F**). Consistent with our expectations from the behavioral dynamics, we show that during trials where the animals respond to salt, neural activity peaks in these regions, and descends to baseline within the trial (**S3D,E**). However, when the animal is unresponsive, the calcium signal plateaus, suggesting that extracting changes in external salt may follow more complicated principles than a simple derivative. Future studies should be able to shed more light on these questions.

### Olfactory input is necessary to drive NaCl avoidance behavior

We next wanted to define the precise roles that the olfactory system and the lateral line each play in the generation of salt avoidance behaviors. To do this, we incubated larvae in various concentrations of copper sulfate, which kills cells in both the lateral line ^27^ and olfactory epithelia ^28^. After treatment and recovery periods, we examined the larvae’s behavioral responses to 50 mM NaCl pulses and found their reaction rate decreased with increasing copper concentration (**4A**). In fact, behavioral responses to NaCl were essentially abolished at a copper sulfate concentration of 20 μM. To ensure that copper does not simply degrade all motor ability, we verified that copper treated larvae showed no reduction in the performance of the optomotor response (**4B**), an innate visuo-motor behavior that leads the animal to follow whole field motion ^29^.

**Figure 4:**
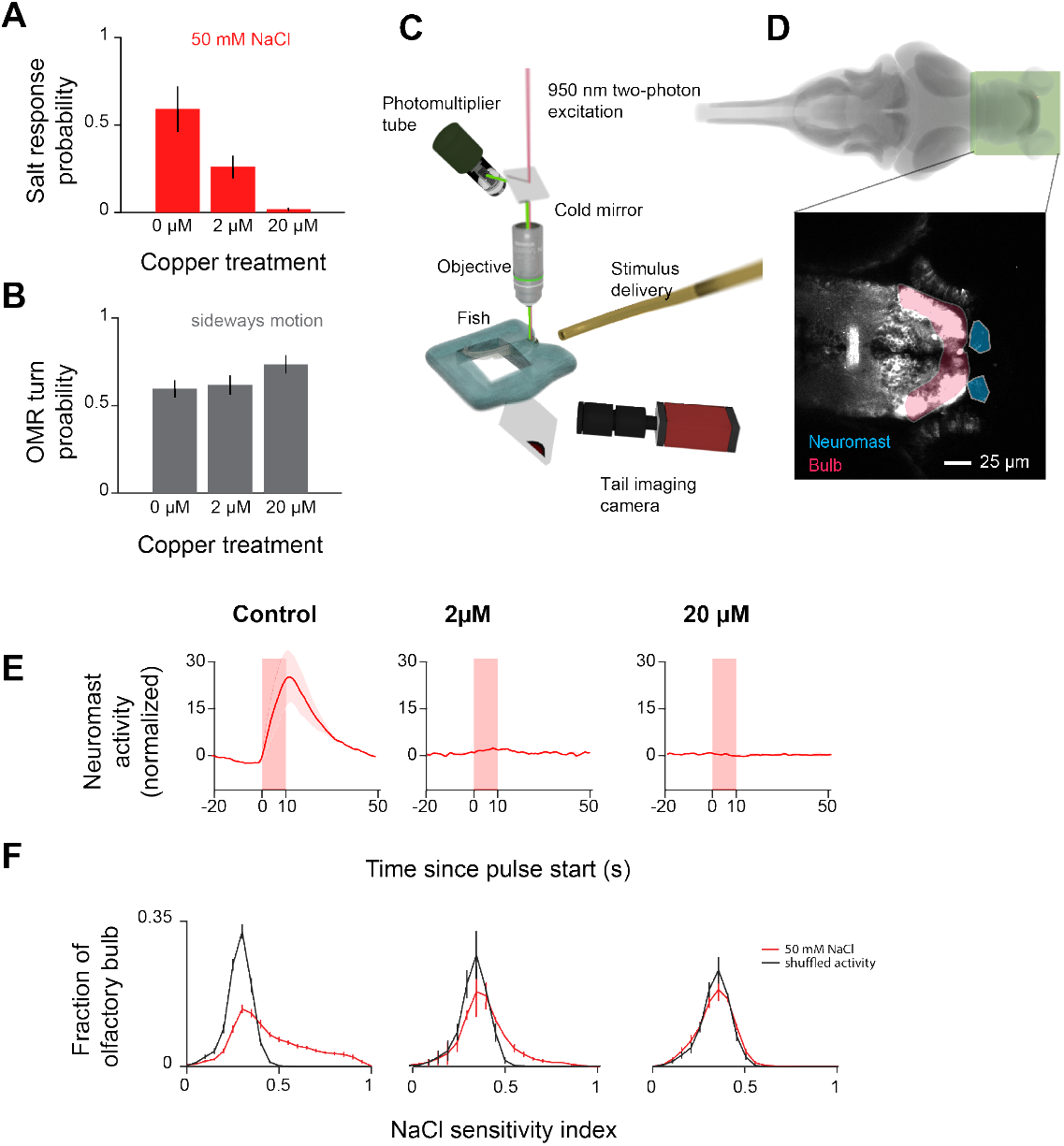
Salt avoidance behavior is abolished after removing olfactory sensitivity to NaCl. **A.** Behavioral response probability of tail embedded larvae to 50 mM NaCl after incubation in different concentrations of copper. **B.** Behavioral response probability of tail embedded larvae to whole field motion after incubation in different concentrations of copper. **C.** Schematic depicting the two-photon microscope setup used to image the olfactory bulb and epithelia. **D.**Region of the brain imaged in this figure. Zoom depicts sample slices averaged over time. Shadings indicate segmented regions - neuromast (orange) and olfactory bulb (red). **E.** Stimulus-triggered averages of neuromast calcium activity during 50 mM NaCl pulses. Each unit is normalized by the variance of its activity during the pre-stimulus period. For fish where neuromasts are fully removed, masks are drawn around where they would normally be. **F.** Distribution of NaCl sensitivity in units across the olfactory bulb where the sensitivity index of a unit is defined as the average correlation of the unit’s calcium response during each trial to the mean stimulus triggered response.

To differentiate the relative importance of olfaction and the lateral line, we examined whether the extent of damage to either modality was predictive of the behavior. To quantify the remaining salt-induced calcium activity in these regions, we wished to have higher spatial resolution that was afforded by the light-sheet microscope, thus we performed 2-photon imaging to assess the extent of the removal of NaCl-sensitivity after copper treatment (**4C,D**). We found that treatment with as little as 2 μM copper sulfate already abolished all lateral line responsiveness, even though the animal continued to respond to NaCl (**4E**, **S4A,B**). By contrast, a significant fraction of NaCl-sensitive olfactory bulb units remained responsive at this concentration (**4F**). Like the behavior, these responses are only completely removed with 20 μM copper sulfate, implicating the olfactory system as critical for NaCl avoidance. To validate the necessity of olfaction for NaCl-triggered behaviors, we next performed a crude yet informative experiment; we rotated the fish 180° relative to the stimulus. This allowed us to expose the neuromasts, as well as other assorted somatosensory systems of the tail to NaCl, while keeping the face and the associated olfactory system largely unaffected. Under this arrangement, fish were significantly less likely to respond than when we subsequently exposed the rostrum of those same fish to NaCl (**S4C**), supporting the notion that olfactory exposure is necessary to elicit NaCl avoidance behavior.

### A sparse subset of olfactory sensory neurons are sensitive to NaCl

We next wished to determine how NaCl-sensitivity might arise in the olfactory system. One possibility is that NaCl directly depolarizes the membranes of all externally contacting neurons, including all olfactory sensory cells. However, we found that fewer than 5 percent of all identified sensory units are responsive to the tested concentrations of NaCl (**5A-C**), suggesting that NaCl activates a specific subpopulation of olfactory sensory neurons. To address the possibility that the large fraction of “silent” neurons may be unresponsive due to some unspecified pathological condition of our preparation, we tested whether these NaCl-insensitive cells might respond to other odorants. To that end we examined two compounds, one noxious to zebrafish, (cadaverine ^30^) the other appetitive (glycine ^31^), and found that both chemicals activated many NaCl-insensitive neurons in the epithelia and bulb (**5D-I**). In fact, while there is some overlap between NaCl and cadaverine sensitive cells, we found zero glycine sensitive cells in the olfactory epithelia that detect NaCl (**5H,I**).

**Figure 5:**
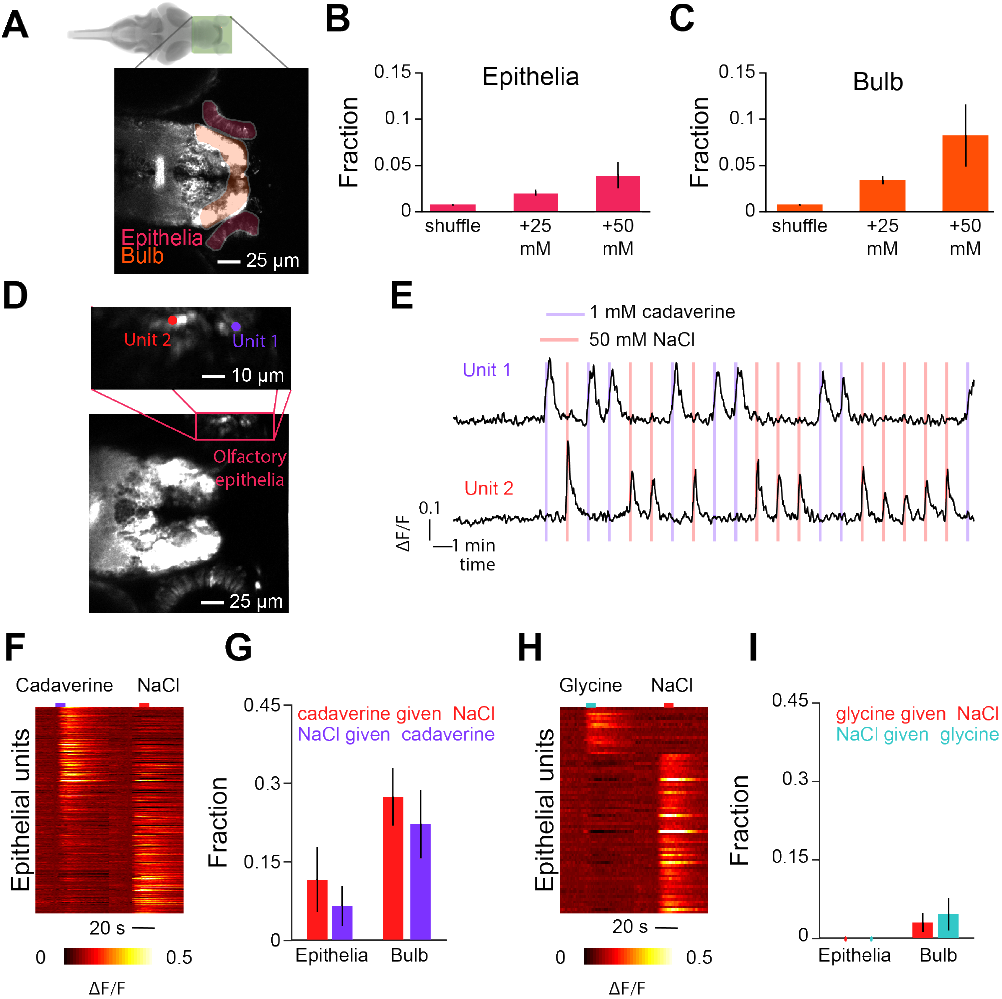
A small fraction of functional olfactory sensory neurons respond to NaCl. **A.** Plane from the imaged region. Shadings indicate segmented regions - olfactory bulb (orange) and epithelia (red). **B.** Average fraction of active units in the olfactory epithelia of *HuC:GCaMP6s* positive fish during 25 mM or 50 mM pulses and after applying the same criteria to shuffled traces (error bars indicate SEM across fish). **C.** Average fraction of active units in the olfactory bulb of *HuC:GCaMP6s* positive fish during 25 mM or 50 mM pulses and after applying the same criteria to shuffled traces (error bars indicate s.e.m. across fish). **D.** Projection across time of a sample slice imaged with 1 mM cadaverine and 50 mM NaCl. Inset depicts the location of two sample units from within the olfactory epithelia. **E.** Calcium traces of the two units depicted in F in response to 10 s pulses of 1 mM cadaverine or 50 mM NaCl. **F.** Heatmap depicting stimulus triggered average activity of all responsive epithelial units to cadaverine and NaCl. **G.** Fraction of units that are responsive to NaCl that are also responsive to cadaverine, and vice-versa (error bars indicate SEM across fish). **H.** Heatmap depicting stimulus triggered average activity of all responsive epithelial units to glycine and NaCl. **I.** Fraction of units that are responsive to NaCl that are also responsive to glycine, and vice-versa (error bars indicate SEM across fish).

Since genetically separable classes of cells detect cadaverine (ciliated) and glycine (microvillous) ^32^, the absence of sensory neurons that detect both glycine and NaCl raised the possibility that only the ciliated class responds to NaCl. However, imaging the NaCl response properties in two transgenic lines that distinguish these classes (*OMP:Gal4/Uas:GCaMP6S* for ciliated and *TRPC2:Gal4/Uas:GCaMP6S* for microvillous), revealed comparable levels of NaCl induced activity (< 5%) in both populations (**S5**), thus allowing us to rule out this hypothesis.

### NaCl-sensitive neurons are driven by sodium and chloride

Having established that sodium chloride does not directly depolarize all epithelial neurons, we next tested whether olfactory responses are driven specifically by sodium and/or chloride or whether they are tuned to an environmental shift that co-occurs with NaCl fluctuation (*i.e.* osmolarity or conductivity). To that end, we presented a given fish with random pulses of either 50 mM NaCl to identify the NaCl-sensitive regions, or a “test” chemical. Recalling the behavioral results in our gradient assay (**1D**), we expect that NaCl-sensitive neurons are not osmolarity sensors, and indeed, we found that NaCl-sensitive cells do not respond to equimolar mannitol (**6A-C**). However, the larvae’s behavioral avoidance of all ionic solutions (**1D**) offers no clues as to how the NaCl-sensitive neurons might respond. On the one hand, these neurons might detect any change in conductivity, while on the other extreme, they could be specifically tuned to sodium or chloride. Thus, we tested a series of solutions to dissect this. When we exposed larvae to KCl, which, in effect, exchanges sodium for a comparable monovalent ion, potassium, NaCl-sensitive neurons were strongly activated (**6A-C**), suggesting they are not sodium-specific. To determine whether chloride drives this activity, we tested sodium conjugated with a different anion, iodide. Yet, this pairing drove the population as well (**6C**). Overall, these results suggest NaCl-sensitive cells at least report the presence of monovalent ions. However, we found they are not conductivity sensors, as exposing the fish to isoelectric or higher concentrations of a divalent cation (magnesium chloride) generates significantly weaker responses in these cells than to NaCl (**6C**, **S6A-D**).

**Figure 6:**
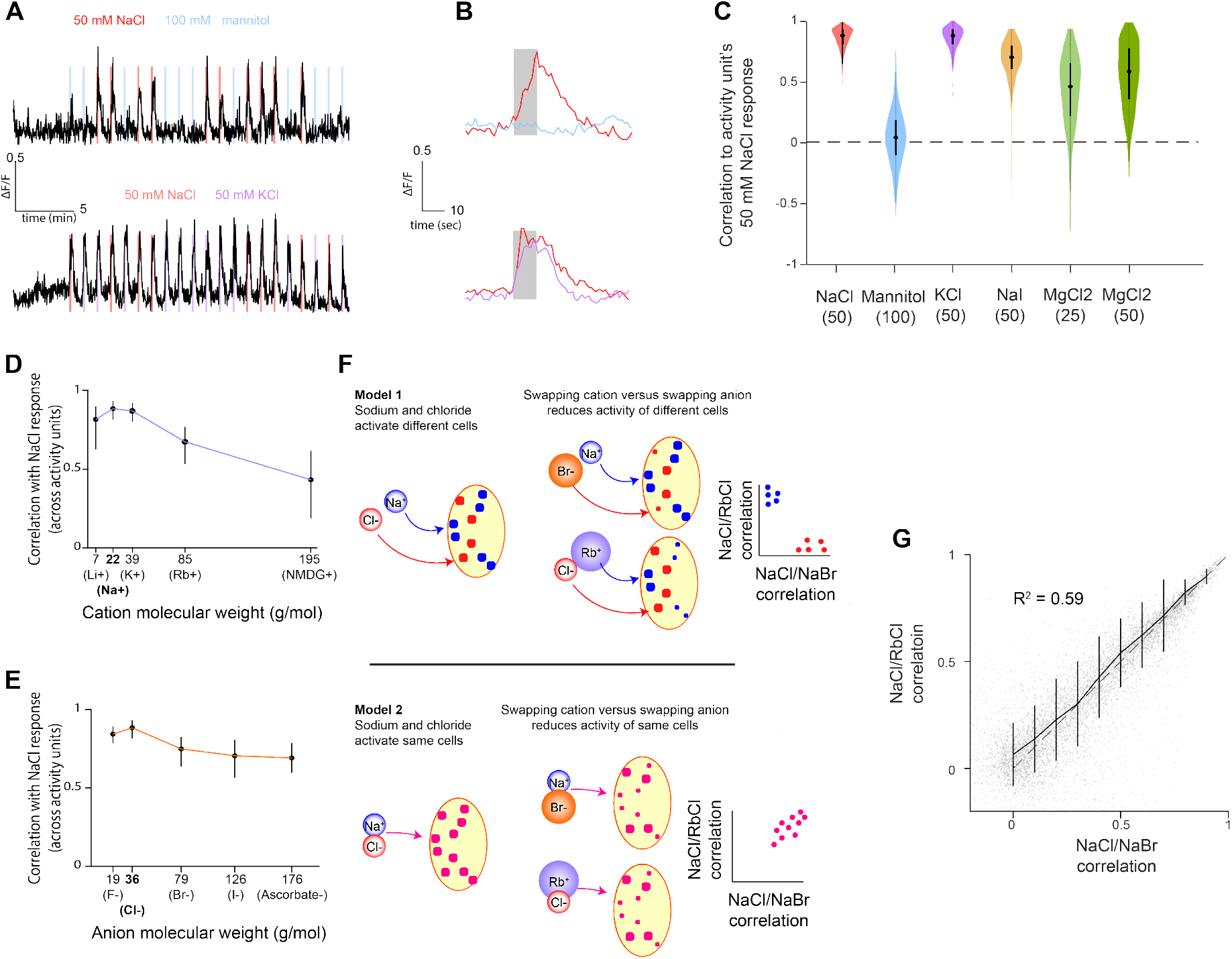
NaCl-sensitive olfactory bulb units are sensitive to sodium and chloride ions. **A.** Sample activity trace from an olfactory bulb unit while the fish is stimulated with pulses of 50 mM NaCl and 100 mM mannitol (top) or 50 mM KCl (bottom). **B.** Stimulus-triggered averages of the activity unit in A in response to 50 mM NaCl and 100 mM mannitol (top) or 50 mM KCl (bottom). **C.** Distribution of the correlation of each activity unit’s NaCl response to different chemicals. Bars in violin plots indicate median ± 25%. **D.** Median (error bars 25-75%) correlation of NaCl activity with different cation/chloride combinations. **E.** Median (error bars 25-75%) correlation of NaCl activity with different anion/sodium combinations. **F.** Cartoon depicting the two models of NaCl sensitivity, and expected results from stimulating NaCl sensitive cells with NaBr and RbCl given each model. **G.** Scatter plot of correlation between NaCl and RbCl responses versus correlation between NaCl and NaBr responses for each activity unit tested (indicated by gray dots, *n*= *3* fish). R-squared calculated from the correlation of all individual units from within the three olfactory bulbs. Solid line indicates average and variance of NaCl/RbCl correlation for binned NaCl/NaBr correlations. Dashed line indicates unity.

Our observation that fish will swim towards deionized water implies that the sensory input driving this behavior should be able to decrease from the baseline established under fish water. To test this, we examined the activity of NaCl-sensitive olfactory neurons exposed to deionized water, and, consistent with the behavior, we observed that their activity decreased (**S6E-G**).

We next wished to determine how different monovalent ions influence the activity of individual NaCl-sensitive units. When we pair chloride with cations larger or smaller than sodium, we find that a greater fraction of olfactory units responds with activity dissimilar to their NaCl response (**6D**). In addition, pairing sodium with different anions generates the same effect (**6E**). This pattern suggests two possibilities: **1)** there exist distinct populations of sodium-sensitive and chloride-sensitive neurons or **2)** there is a single population of cells equally sensitive to both sodium and chloride (**6F**). To distinguish between these two hypotheses, we compared the activity induced by RbCl and NaBr, two ion-pairs that have similar effects at the population level. With these two solutions, we asked whether swapping either sodium or chloride reduced the activity of distinct sets of neurons (**6F**). We found, instead, a strong correlation between the activation of each individual neuron by both RbCl and NaBr (**6G**). Following this result, we propose that NaCl-sensitive neurons in the zebrafish are tuned to the presence of both sodium and chloride, thereby implicating a novel and undescribed molecular mechanism for environmental salt detection.

## Discussion

Salt detection is classically considered a task for the gustatory system. This view reflects a focus on understanding salt sensing in terrestrial animals ^5,7,33^. On land, animals interact with environmental salt through ingestion, making taste the primary conduit for salt-related decisions. For teleosts and other aquatic organisms, however, saltiness is a critical environmental variable, which critically restricts their possible habitats. Here, we have discovered that at least one species of fish uses olfaction to detect salt. Unlike taste, which in teleosts is reportedly exclusively used for ingestion related decisions ^34^, olfaction has broad influence ^35^, being involved in behaviors as diverse as finding food ^36^, mating ^37^, and avoiding dangerous environments ^30,38^. Our finding thus suggests interesting implications for the evolution of cross-modal behaviors. Rather than endow a salt-sensitive modality, such as taste, with the ability to regulate navigational behaviors, natural selection has led to the incorporation of salt sensitivity into a different modality, olfaction, that already influences navigation across animals as diverse and flies and mice ^39,40,41^.

Further, when we investigated the detailed dynamics of the salt gradient navigation, we found that larval zebrafish modify their behavior during the onset of increased salinity (Figure 1N). Yet, the heightened activity in the olfactory system is sustained throughout the duration of elevated salt. This raises several questions. First, how does the larva’s brain generate the relevant derivative? At present, the nature of this computation is unclear. While the interpeduncular nucleus as well as the raphe seem like potential sites to explore this further, their activity profiles do not reflect a pure derivative of salt concentration. In particular, the lack of neural adaptation during trials without a behavioral response obfuscates the possible neural implementation. Future studies should further dissect the nature of the behavioral algorithm by which changes in salt concentration are detected, as this may offer clear constraints to the neural mechanisms. For example, feedback from an efferent relay of the animal’s motor activity may modulate how the animal’s brain processes sustained salt elevation. Second, why does the fish expend extra energy to sustain high firing rates in the olfactory system when the behaviorally relevant information only lasts a few seconds? Even the baseline activity in normal fish water is well above the minimal activity during exposure to deionized water. In line with a hypothesis discussed in Lovett-Baron et al.^19^, retention of information about the surrounding salinity might be necessary for regulating non-motor related functions. After early attempts to escape salt fail, the animal may still survive if it can hormonally regulate its ion balance. For example, salt information from the olfactory system may directly regulate hypothalamic release of prolactin ^42^ or cortisol ^43^ to balance ion uptake and excretion, respectively.

At present, the molecular mechanisms that endow hair cells and some olfactory sensory neurons with NaCl-sensitivity are unclear. Here, we observe that zebrafish larvae avoid levels of salt concentrations that fall within the operating range of the epithelial sodium channels that are essential for detecting appetitive salt in land animals ^5^. Yet, fish lack these channels, suggesting they would need to utilize some other mechanism. Indeed, the response profile of the larvae’s NaCl-sensitive olfactory neurons suggests a different mechanism. Instead of being specifically activated by sodium like the terrestrial sodium channels, these cells are broadly activated by both monovalent cations and anions. This response profile is also distinct from known molecules responsible for detecting internal sodium, which are either specific to high (>100 mM) concentrations of sodium ^44^ or are general osmolarity sensors ^45^. Whether the cellular responses result from the action of multiple receptors and channels, or from a single protein, i.e. an ion-pair receptor ^46^ is unclear.

To our knowledge, the receptor that most closely matches the ion-sensitivity profile of the larvae’s olfactory sensory neurons is *pickpocket23* in the fruit fly ^47^. Like the zebrafish cells, this protein is broadly sensitive to monovalent ions, and not osmolarity. However, the relative influence of the cation and anion is undescribed. Further, much higher concentrations (>200 mM) are needed to drive activity in *pickpocket23* positive cells than in the zebrafish sensory neurons, suggesting, at a minimum, that zebrafish use alternative secondary mechanisms. Identifying the responsible molecules should be the work of future studies. The robust nature of the avoidance behavior, as well as the ability to test many larvae simultaneously in our lane assay, make this question well-suited for a broad genetic screen.

## Methods and Materials

### Animal Husbandry

Unless otherwise noted, all fish used were the offspring of crosses of *HuC:GCaMP6s* positive and *nacre* +/− parents. Embryos were raised at 27 degrees celsius. For the first 24 hours embryos developed in embryo water plus methylene blue. Afterward, larvae were exclusively raised in filtered (200 nm pore size) facility water. Water was exchanged every day. The larvae were fed live paramecia starting at 4 days post-fertilization (dpf). Experiments were performed on fish between 6 and 7 days old, unless otherwise noted. All experiments followed institution IACUC protocols.

### Free-swimming place-preference assay

In order to test whether larval zebrafish avoid salts, we sought to develop a rig that would allow us to determine a larva’s place preference within a chemical gradient. Previous studies have examined chemical preference in adult zebrafish by using flow chambers. These strategies, however, introduce two main confounds that we wished to avoid. First, any chemical avoidance behaviors will be convolved with rheotaxis and the optomotor response, and switching chambers forces the larva to behave in opposition to its rheotactic drive. Second, these arenas only create a very steep local gradient at the chamber boundaries, which may not provide the information necessary for the larva to find its preferred location.

Agar pads are made from 3% low melting point agarose. Agar is poured into premade casts designed to fit the arena. After the agar settles, it is cut out and added to the arena, which is then filled with water, lane by lane. We confirmed the presence of a gradient by using a refractometer to measure the osmolarity of the water in 15-minute intervals at the quartiles of the arena’s length. After the fish are added the initial background image is calculated for 20 seconds and the experiment begins. To make ionic solutions, salt (all purchased from *Sigma-aldrich*) was added to the agarose prior to melting.

### Behavior analyses

All analyses were performed with custom-written MATLAB code. Preference indices (P.I.) were calculated based upon the position of the fish along the axis perpendicular to the two agar pads. We define a preference index as the difference in time spent on the half of the arena close to the “test” pad from the time spent away divided by the total length of the experiment:

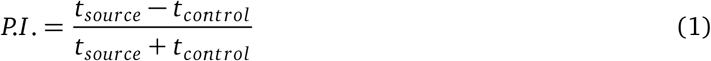

where *t*_*source*_ and *t*_*control*_ are the time spent in the half of the arena closest to the salt and control, respectively. As such, the preference index ranges from one for larvae that spend all of their time on the side proximal to the NaCl, and negative one for those that spend all of their time distal to the salt. To analyze kinematic parameters, bouts were segmented automatically from the absolute speed of the fish combined with identifying periods of high variance in the heading angle. Bouts were then separated by whether they brought the animal closer to or further from the source agarose pad (by at least 0.2 mm).

### Free swimming simulations

For simulations, we assumed larvae chose bouts from one of two types of swim events - straight swims and turns. We described the heading angle changed during these swims by a gaussian, 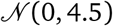, and lognormal, 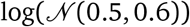, distribution for straight swims and turns, respectively. For every swim, larvae choose a heading angle turn from a probability distribution that is a linear combination of the above distributions. As the virtual fish navigates, it experiences changes in salt concentrations. We simplify the gradient by estimating it as a linear increase from the control agar pad to the source that rises linearly with time. We fit slopes to reach a peak concentration of 15% of the source at the end of the experiment. Following each bout, we redraw the heading distribution, based upon the change in salt concentration. As the change in salt increases, we apply a higher weight to the turn distribution. This weight is assumed to be a linear function of the change in salt concentration caused by the previous bout. Bouts are assumed to take place at a frequency of 0.8 Hz and move the fish by 1 centimeter.

### Head-embedded chemical stimulation

At present, the designs available for performing calcium imaging of the brains of freely-swimming zebrafish larvae ^48^ do not offer the same resolution as generated by traditional methods that require the brain to be immobilized, such as two-photon point scanning and light-sheet microscopy. Therefore, we designed a preparation that would enable simultaneous stimulation of a head-immobilized larva with salt and recording of its behavior. In this preparation, we use gravity to control fluid flow that is directed to the rostral end of the fish by a narrow, 360 μm diameter perfusion pencil tip (*AutoMate Scientific* 04-360). The flow speed was approximately 1.5 ml/minute. Multiple solutions were passed through the perfusion tip via an 8-channel manifold that ensured rapid liquid volume exchange (*AutoMate Scientifi*c 04-08-zdv). Solution outputs were regulated by solenoids (*Cole Palmer EW-01540-01*) via an Arduino^®^ during the experiment such that at all times one and only one solution was being presented to the fish. Presenting a continuous stream of flow both attenuates behavioral responses to sudden changes of flow velocity and hastens the removal of salt at the end of a trial compared to diffusion alone. The dynamics of the pulse were assessed by imaging pulses of 10 nM fluorescein under an epifluorescent microscope (Olympus^®^ MVX10).

### Copper treatments

For olfactory and lateral line ablations, fish were treated with either 0, 2, or 20 μM Copper Sulfate (CuSO_4_). Fish were incubated for one hour in the chemical and then given one hour to recover before behavioral experiments. Ablation of the lateral line was confirmed anatomically by incubating the fish in a dye, FM 1-43 (*ThermoFisher* T3163) for five minutes followed by a 15-minute wash in fish water and taking images under an epifluorescent microscope.

### Light-sheet microscopy

Volumetric imaging experiments were performed with a custom-built single-photon lightsheet microscope similar to that described previously ^49^. One difference, however, is that we used a transparent specimen chamber and holder to enable tail tracking via a camera below the fish. For imaging, *Nacre* −/− larvae positive for *GCaMP6s* expression under the *HuC* promoter ^48^ were embedded in agarose. To stimulate the fish, we removed the agarose surrounding the nose. To allow behavioral monitoring, we also removed the agarose that surrounds the tail. Larvae were illuminated with a 488 nm digitally scanned sheet that swept through 200 μm of depth with 4 μm steps at 1 Hz. During the experiment, fish were stimulated with 10 s of a given concentration of NaCl (25, 50, 75, 100 mM) separated by 40s of water flow. Each concentration block lasted for 5 minutes and was separated by 5 minutes.

### Two-photon microscopy

Two-photon microscopy experiments were performed with a custom-built two-photon microscope described previously ^50^. *Nacre* −/− larvae positive for *GCaMP6s* expression in the brain were embedded in 1.8% agarose, and the tail and nose were freed as done during light-sheet experiments and embedded behavior experiments. Larvae were imaged with a spectra-physics *Mai-tai* laser at 950 nm with 10 mW of power at the sample. Volumes spanning the olfactory bulb were imaged plane-by-plane at 8 μm steps.

### Image Analysis

After imaging experiments, all data was segmented into activity units that approximate cells. Segmentation was performed using an algorithm based on the one developed by Portugues et al.^23^. The eyes, which are heavily autofluorescent, were manually masked out of each stack to avoid segmentation and registration errors. Registration to the reference brain was performed using the Computational Morphometry Toolkit () ^51^ as described previously ^24^. The following parameter set was used:

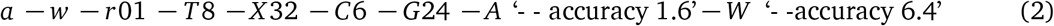

The relationship between the stimulus and each activity unit was determined by calculating their mutual information. To do so, each neural signal was reduced to a 300 block time series, where each block was the average activity from 10 seconds. This made the score blind to dynamics within the pulse, so, for instance, stereotypic differentiating cells could also be discovered. The resulting signal was normalized to range from 0 to 1 and then binned into 10 equal-sized units. The same was done with the stimulus signal. A behavioral signal was generated by counting the number of behavioral responses in each 10-second block. A combined behavior and stimulus signal was generated by multiplying the behavior and stimulus signals. Mutual information was then defined as follows:

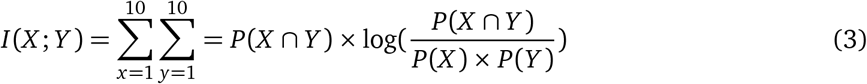

Here, x and y represent stimulus and activity bins. Scores were then normalized by the entropy of the stimulus.

To test the classification ability of given regions, support vector machines were trained on a random subset of 50% of trials to identify from the population activity within that region either the concentration of salt in the trial, or whether the animal responded during the trial. The resulting classifier was then cross-validated with the remaining trials. We report the performance of this cross-validation. For each fish, this process was repeated 50 times and performance was averaged across these repetitions.

For two-photon imaging experiments, we defined units as active based upon their coherence across trials. Namely, we asked for each trial, what is the correlation of the units activity with the average across trials. All units with an average correlation across trials greater than 0.7 were deemed active. This threshold was determined by the value above which the probability of being seen in shuffled data is less than 0.01. For comparisons across fish, calcium traces were normalized by the variance of the unstimulated activity.

### Data Availability

Analysis code and data will be made available upon final acceptance of the manuscript.

## Author Contributions

KJH conceived the project with FE and performed all experiments. DGN and TP built the light-sheet and two-photon microscopes. KJH analyzed the data. KJH wrote the manuscript with input from FE and DGN.

## Acknowledgements

KJH received funding from the Harvard Minds Brains and Behavior Initiative. NIH. We thank all members of the Engert lab for support and advice throughout the project. FE received funding from the National Institutes of Health (U19NS104653, R43OD024879, 2R44OD024879), the National Science Foundation (IIS-1912293) and the Simons Foundation (SCGB 542973).We also thank Mariela Petkova and Hanna Zwaka for providing valuable feedback and suggestions to improve the manuscript.

---

**Figure S1:**
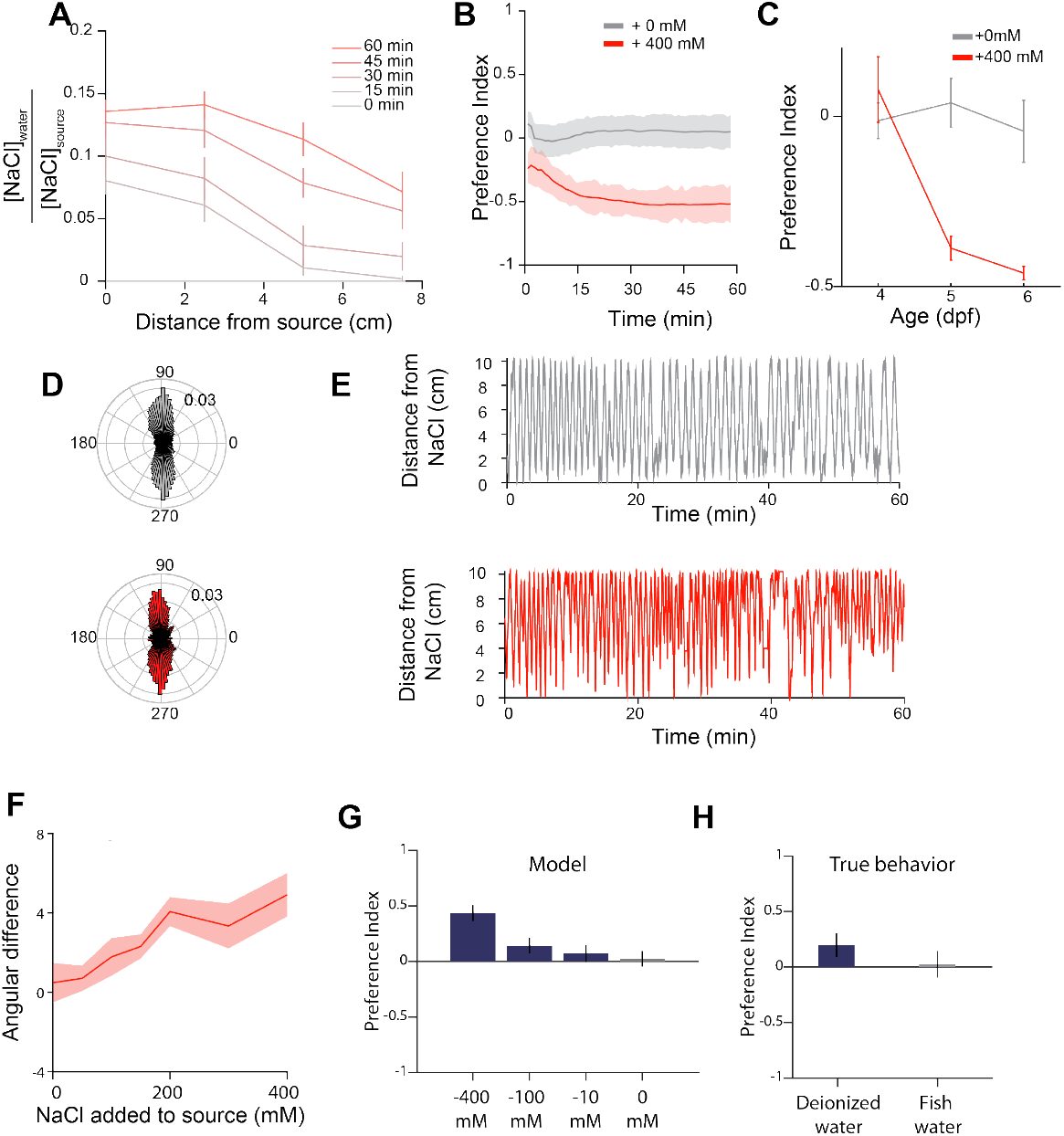
**A.** Development of the salt gradient over time relative to the concentration of salt added to the source. **B.** Evolution of preference indices over the course of the experiment. **C.** Avoidance behavior by larval age. **D.** Polar histogram of orientations of fish in no gradient (top) and a gradient generated from 400 mM NaCl (bottom). Fish facing 270 degrees are facing the source. **E.** Sample positional trace of a fish experiencing no gradient (top) and a gradient generated from 400 mM NaCl (bottom) along the axis that bisects the two agar pads. **F.** Difference in average angle for fish climbing the gradient versus descending for different concentrations of source NaCl. **G.** Model predictions for preference index of larvae navigating “negative” salt gradients. **H.** Actual preference index of a fish navigating toward deionized water in the source agar pad.

**Figure S2:**
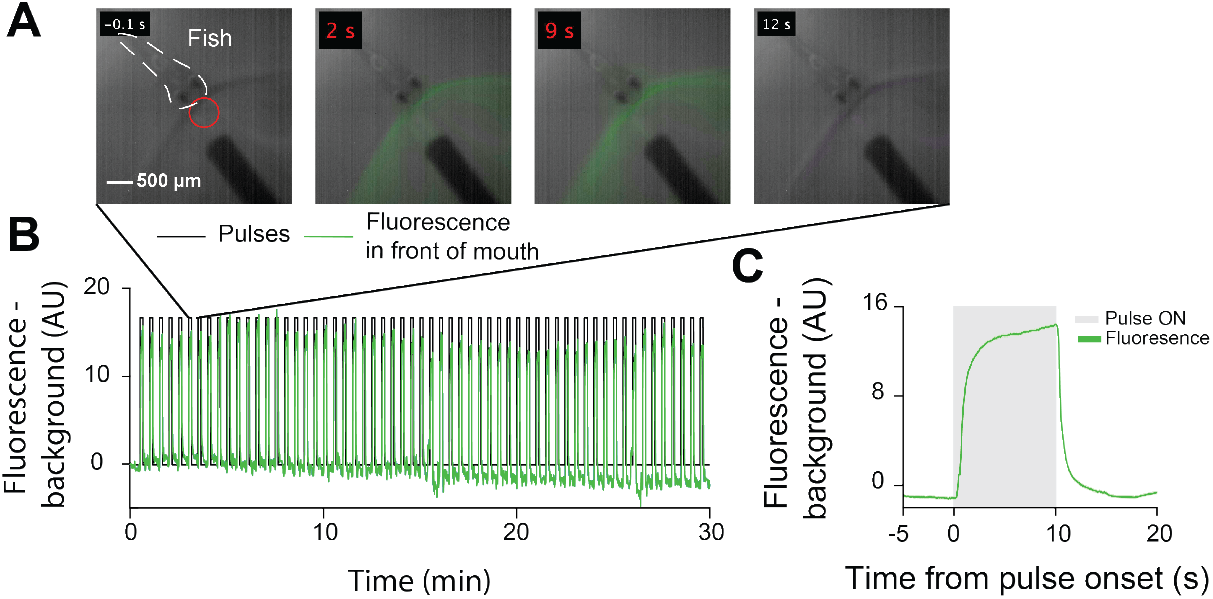
**A.** Example frames of a ten-second 10 nM fluorescein pulse. Red circle indicates the region analyzed in **B** and **C.** Dashed white outline surrounds the fish. Time is relative to the beginning of the fluorescein pulse, and red text indicates presentation of fluorescein. **B.** Fluorescence over the course of thirty minutes with 30 second interstimulus intervals. **C.** Average fluorescence changes during a pulse.

**Figure S3:**
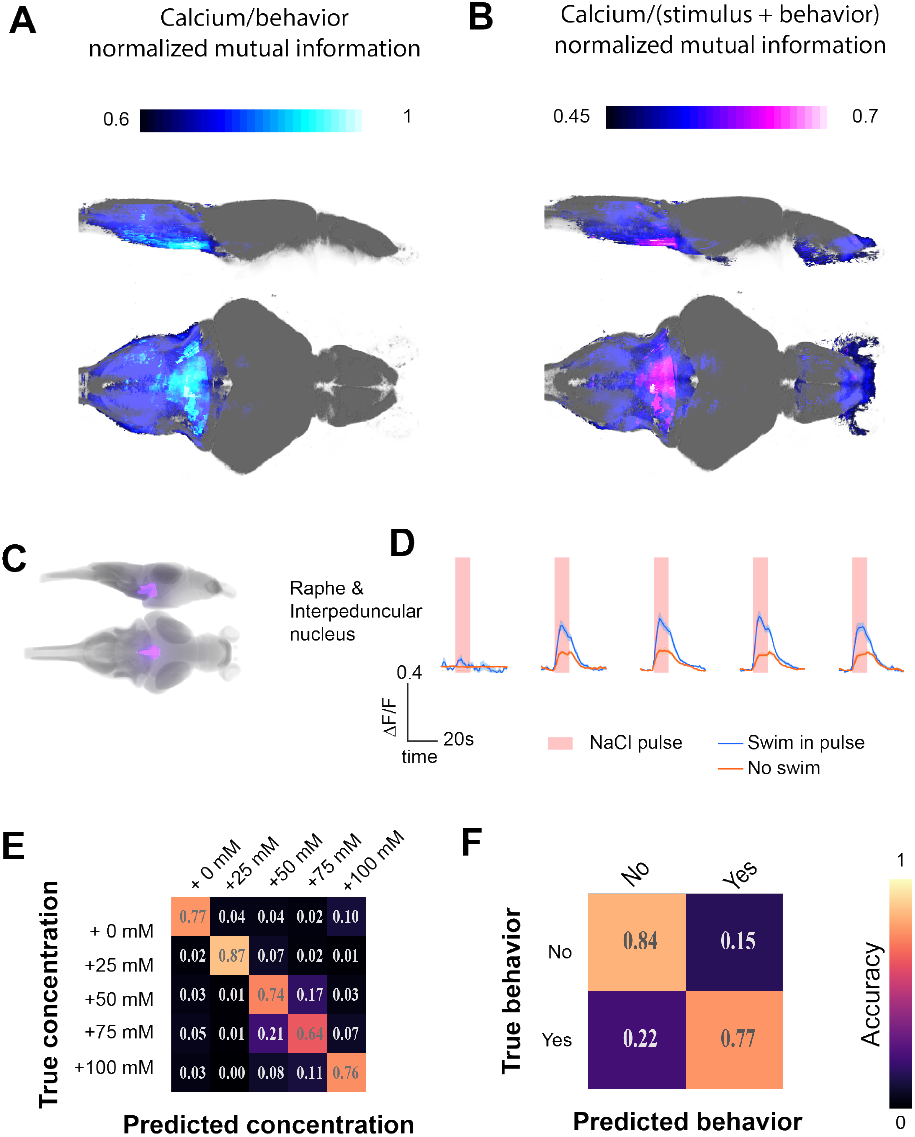
**A.** Map of mutual information with behavioral responses across all fish. **B.** Map of mutual information with NaCl concentration and behavioral responses across all fish. **C.** Location of the Raphe and interpeduncular nucleus masks within the *Z-brain*. **D.** Average responses from these regions across fish during trials where the fish attempted to swim away and those where they did not. **E.** Confusion matrix depicting *stimulus* classification accuracy of support vector machines trained from activity in the raphe and interpeduncular nucleus. **F.** Confusion matrix depicting *behavioral response* classification accuracy of support vector machines trained from the dorsal raphe and interpeduncular nucleus.

**Figure S4:**
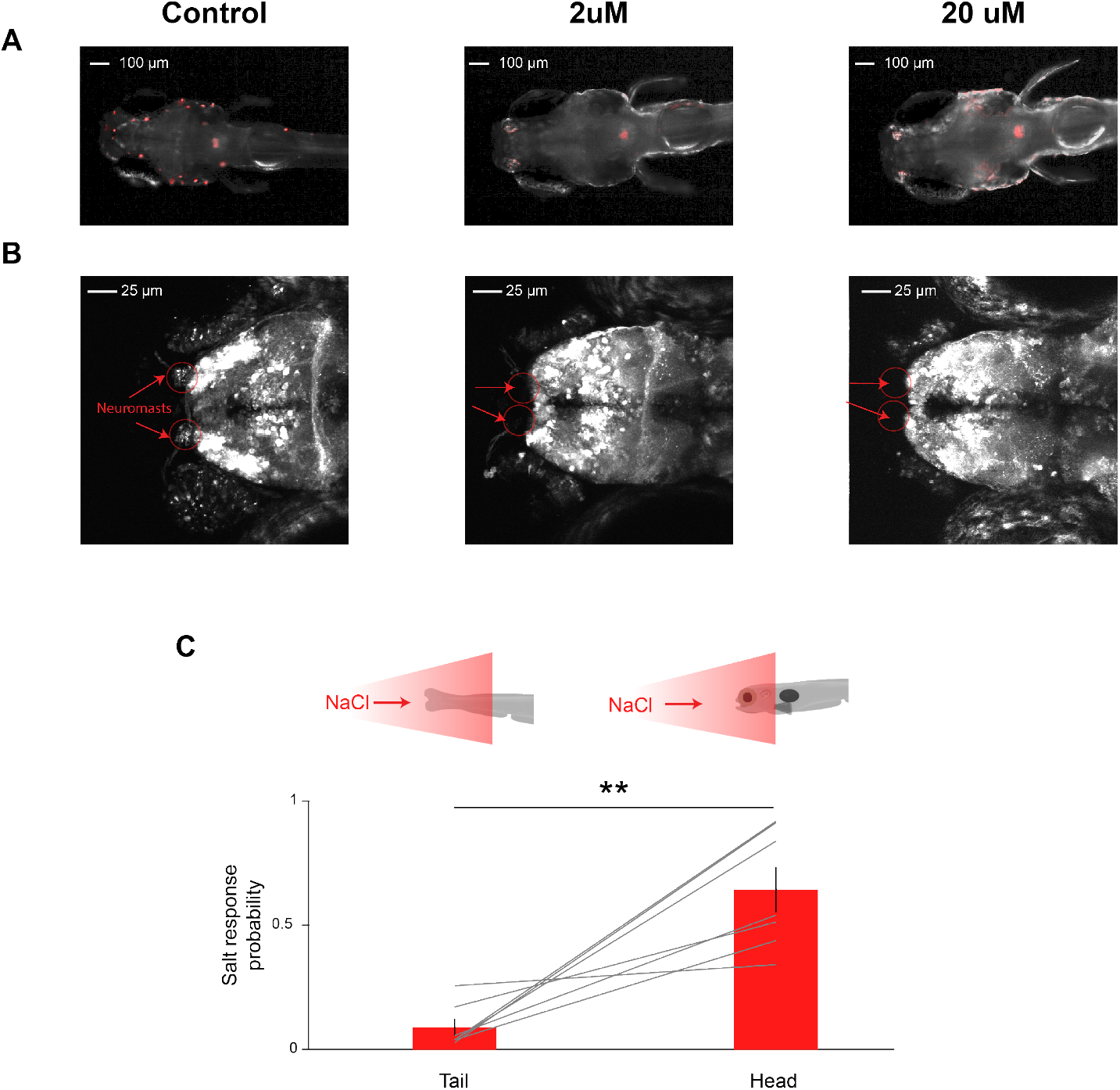
**A.** Epifluorescence images of fish stained with FM1-43 to label neuromasts (red) **B.** Maximum intensity projections of the average fluorescence during 2-photon imaging experiments following treatment with different concentrations of copper sulfate. Red arrows indicate where the nasal neuromasts should be located. **C.** Behavioral response rate to 50 mM NaCl if the stimulus is presented to the tail or to the head (Wilcoxon signed-rank test, *p* < *0.01*). Gray lines indicate individual fish. Tail experiment always preceded head experiment.

**Figure S5:**
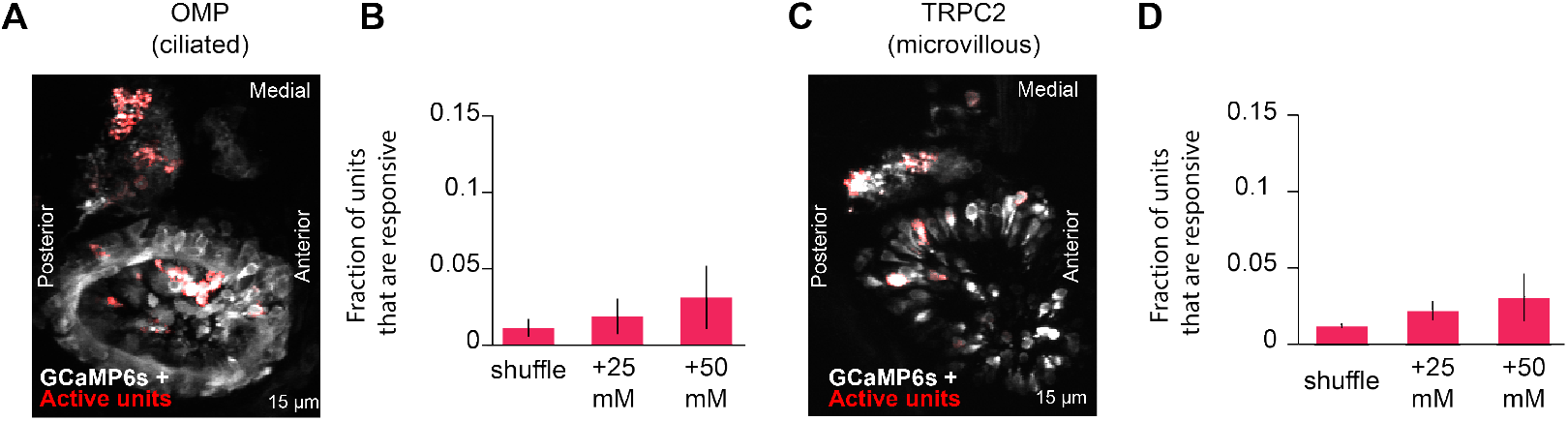
Hunger state modulates eye position and bout-type selection. **A.** Maximum intensity projection of two-photon stack of *OMP:Gal4/UAS:GCaMP6S* positive fish with location of NaCl responsive units overlaid (red). **B.** Average fraction of activity units in the olfactory epithelia of *OMP:Gal4/UAS:GCaMP6S* positive fish that were deemed to be active during 25 mM or 50 mM NaCl pulses and after applying the same criteria to shuffled traces (error bars indicate SEM across fish). **C.** Maximum intensity projection of two-photon stack of *TRPC2:Gal4/UAS:GCaMP6S* positive fish with location of NaCl responsive units overlaid (red). **D.** Average fraction of activity units in the olfactory epithelia of *TRPC2:Gal4/UAS:GCaMP6S* positive fish that were deemed to be active during 25 mM or 50 mM NaCl pulses and after applying the same criteria to shuffled traces (error bars indicate SEM across fish).

**Figure S6:**
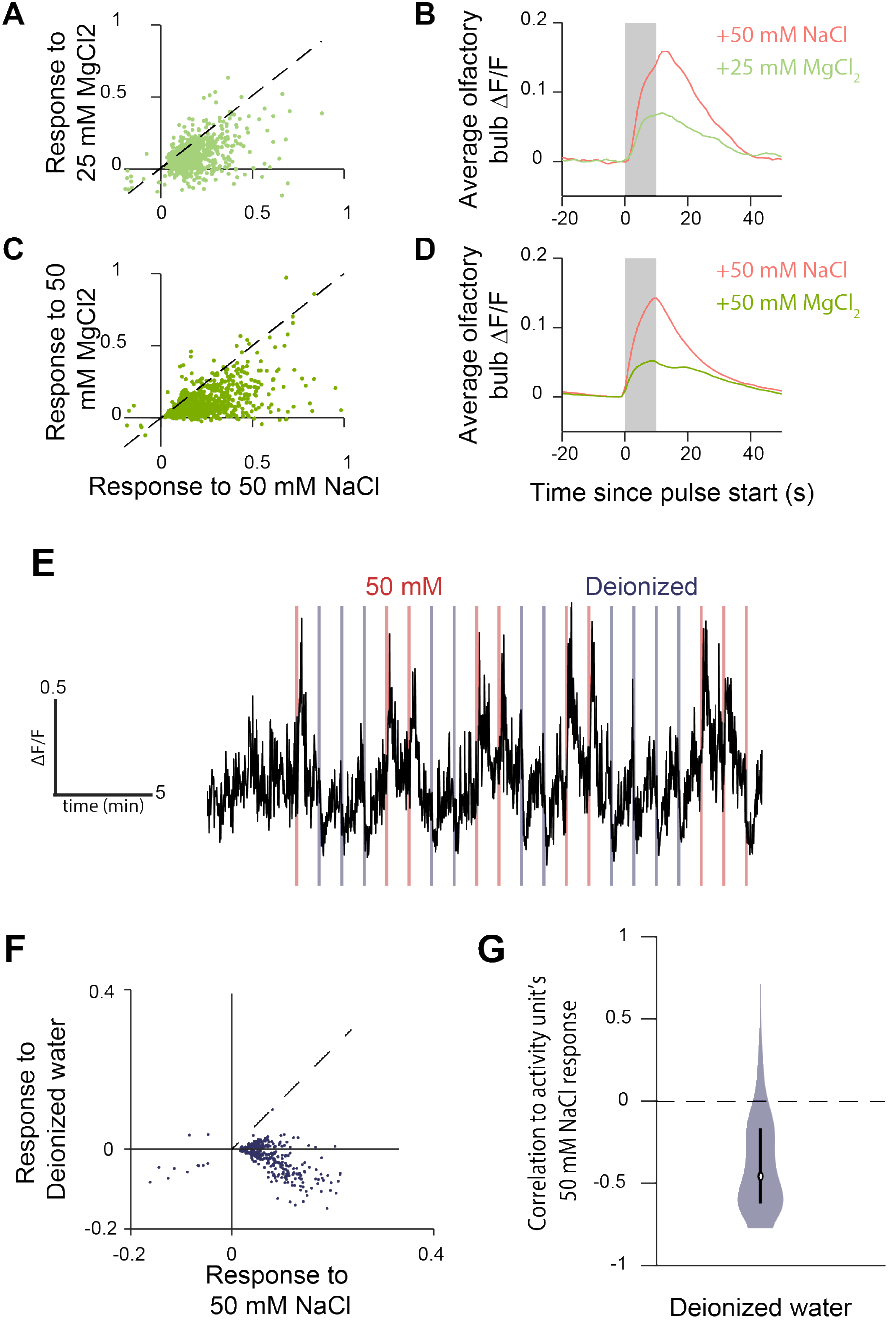
Subdividing bout-classes by kinematic parameters to get bout-types. **A.** Scatter plot of unit’s response to 50 mM NaCl versus 25 mM MgCl_2_. **B.** Average activity induced across the olfactory bulb by 50 mM NaCl and 25 mM MgCl_2_. **C.** Scatter plot of unit’s response to 50 mM NaCl versus 50 mM MgCl_2_. **D.** Average activity induced across the olfactory bulb by 50 mM NaCl and 50 mM MgCl_2_. **E.** Sample calcium trace from a fish stimulated with pulses of 50 mM NaCl and deionized water. **F.** Scatter plot of each unit’s peak calcium response to NaCl and to deionized water response. Each circle represents one unit. G. Violin plot that illustrates distribution of the correlation of a unit’s calcium signal in response to NaCl with it’s response to deionized water. Bars in violin plots indicate median ± 25%.

## Supplementary Video Legends

**Figure SV1: Animation of brain-wide imaging preparation.**

